# Network-Guided Sparse Subspace Clustering on Single-Cell Data

**DOI:** 10.1101/2022.12.20.521229

**Authors:** Chenyang Yuan, Shunzhou Jiang

**Affiliations:** Department of Biostatistics and Bioinformatics, Rollins School of Public Health, Emory University, Atlanta, GA 30322, USA; Department of Biostatistics, Epidemiology and Informatics, Perelman School of Medicine, University of Pennsylvania, Philadelphia, PA 19104, USA

**Author notes:** Correspondence: Chenyang Yuan.

## Abstract

With the rapid development of single-cell RNA sequencing (scRNA-seq) technology, people are able to investigate gene expression at the individual cell level. Identification of cell types via unsupervised clustering is one of the fundamental issues in analyzing single-cell data. However, due to the high dimensionality of expression profiles, traditional clustering methods are difficult to generate satisfactory results. To address this problem, we designed NetworkSSC, a network-guided sparse subspace clustering (SSC) approach. NetworkSSC is based on a similar assumption in SSC that the expression of cells within the same type lies in the same subspace. Moreover, it integrates an additional regularization term to include the gene network’s Laplacian matrix, so as to utilize the network information. The comparison results of 5 scRNA-seq data sets show that NetworkSSC outperforms ordinary SSC and other clustering methods in most cases.

## 1 Introduction

Single-cell RNA-sequencing (scRNA-seq) is a powerful technology that enables the measurement of transcriptomic profiles at the individual cell level (Tang et al., 2009). It achieves unprecedented resolution for the study of transcriptome and provides critical insights into the genomic heterogeneity among individual cells (Buettner et al., 2015).

One of the most common practices in scRNA-seq data analysis is cell clustering (Kiselev et al., 2019), which groups a population of cells based on their transcriptomic similarity, and then labels the groups by cell types or subtypes (Wei et al., 2022). Cell clustering allows many downstream analyses to be carried out, including cellular composition estimation and rare cell type discovery (Chen et al., 2019), and hence is a fundamental step performed in scRNA-seq data analysis. However, the expression profiles often have high dimensionality, i.e., the number of features (genes) is often much larger than the number of samples (cells), leading to statistical challenges in comparing cells in such high-dimensional spaces (Andrews and Hemberg, 2018).

There are various categories of clustering methods applied to scRNA-seq data. Traditional methods include K-Means (Hartigan and Wong, 1979) and spectral clustering (Ng et al., 2001). K-Means aims to minimize within-cluster distances. Spectral clustering clusters the eigenvectors of the graph Laplacian matrix of the data and often outperforms K-Means in real applications. Due to the strong capability of representation learning, deep neural networks have also been applied to clustering in recent years. For instance, DEC (Xie et al., 2016) learns a mapping from the data space to a lower-dimensional feature space where it iteratively optimizes a clustering objective. scDeepCluster (Tian et al., 2019) learns a latent embedded representation optimized for clustering high-dimensional input in a non-linear manner. DESC (Li et al., 2020) learns a low-dimensional representation of the data and adds an encoder to the neural network to cluster cells iteratively. Despite the success of deep learning clustering methods, their performance heavily relies on the neural network structure, hyperparameters, and optimization algorithm, which are difficult to determine and often require strong domain experience. In addition, the interpretability of deep learning clustering methods is much lower than K-Means and spectral clustering. Indeed, for scRNA-seq data analysis, we require accurate, convenient, and stable clustering algorithms with high interpretability.

Sparse subspace clustering (SSC, Elhamifar and Vidal, 2013) is a variant of spectral clustering and has shown promising performance in clustering high-dimensional data. SSC assumes that the high-dimensional data are distributed in a union of multiple low-dimensional subspaces, and each data point can be represented by other data points in the same subspace. SinNLRR (Zheng et al., 2019) further justifies the application of SSC in cell clustering by considering the subspace characteristics of cells’ expression. It assumes that the expression of cells within the same type should lie in the same subspace. Based on the same assumption, AdaptiveSSC (Zheng et al., 2020) modifies SSC and realizes cell type identification by using a data-driven adaptive sparse constraint to construct the similarity matrix. Still, these SSC-based methods also have certain limitations. In cell clustering, it is possible that most of the genes are noisy and not discriminative, which leads to low clustering accuracy, although a few researchers have proposed to use principal component analysis (PCA) to reduce the dimension of the data (Usoskin et al., 2015; Yau et al., 2016). SSC and its variants, however, lack an automatic and effective feature selection procedure to denoise the data. On the other hand, gene co-expression network is very useful prior knowledge in capturing the data information, which is not considered in SSC and its variants.

In this paper, we propose a network-guided sparse subspace clustering method called NetworkSSC. NetworkSSC follows the similar subspace assumption in SSC and includes the gene network’s Laplacian matrix as a regularization term. This method effectively utilizes the gene co-expression relationship from the network structure and is more applicable to clustering gene expression data. It obtains an improved performance on multiple real data sets compared with traditional SSC.

The rest of the paper is organized as follows. Section 2 introduces the idea of traditional SSC, which serves as the basis of our proposed method. Section 3 introduces the design of NetworkSSC and presents the detailed procedures to solve the novel optimization problem. In Section 4, we apply NetworkSSC to five real data sets and compare its performance with five competitive methods. Section 5 closes the paper with a discussion outlining the potential improvements of NetworkSSC.

## 2 Sparse Subspace Clustering

High dimensionality often causes problems to data processing and analysis (Donoho et al., 2000), and SSC uses subspace representation to simplify the clustering process on high-dimensional data. In the subspace assumption, high-dimensional data are distributed in the union of multiple low-dimensional subspaces (Parsons et al., 2004), and a sample can be represented as the linear combination of other samples in the same subspace.

Based on the subspace assumption, the expression of a cell can be written as the linear combination of other cells. In specific, if we denote the expression matrix as *X* ∈ ℝ^*p×n*^, where each row represents a gene and each column represents a cell, then the expression vector for cell *i* can be written as

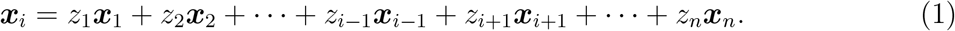

For *j* ≠ *i*, *z_j_* ≥ 0 denotes the similarity between cell *i* and *j*, where *z_j_* = 0 indicates that cell *i* and cell *j* are not of the same type. Extending this expression to all cells results in

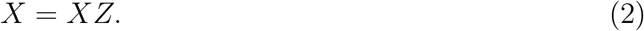

Here *Z* ∈ ℝ^*n×n*^ is the coefficient matrix, and *z_ij_* is the similarity score between cell *i* and *j*. To ensure the properties of *z_ij_*’s, one can impose sparsity constraints on *Z* and force it to have a block diagonal structure, which eventually reveals the subspace structure of the data. The sparsity of a vector is captured by its *ℓ*_0_ norm, i.e., the number of non-zero elements. In real applications, however, the optimization problem containing *ℓ*_0_ norm is often NP-hard, and people introduce *ℓ*_1_ norm as a relaxation of *ℓ*_0_ norm (Chen et al., 2001). In this case, the coefficient matrix *Z* can be computed by

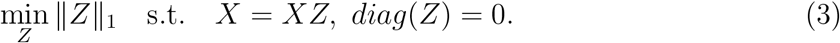

The constraint *diag*(*Z*) = 0 is designed to avoid the trivial solution, where each data is only represented by itself, and *Z* becomes the identity matrix. After relaxation, the optimization problem in (3) can be re-written as

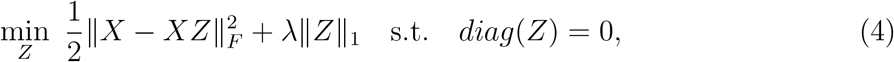

where parameter *λ* is introduced to control the sparsity of the coefficient matrix *Z*.

After solving the optimal *Z*, a symmetric affinity matrix *W* ∈ ℝ^*n×n*^ is constructed by

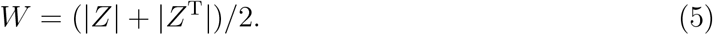

Finally, spectral clustering is performed on the learned affinity matrix *W*, and SSC obtains the clustering result through this final step.

## 3 Network-Guided Sparse Subspace Clustering

We adapt a sparse subspace clustering method tailored to clustering gene expression data, termed NetworkSSC. NetworkSSC is based on the similar subspace assumption in SSC that the expression of cells within the same type lies in the same subspace. It further utilizes the information from the gene co-expression network to exploit the intrinsic relationship within the data. The pipeline of NetworkSSC is outlined in Figure 1. Given an input expression matrix *X*, NetworkSSC constructs the corresponding gene network and Laplacian matrix *L*. It then performs dimension reduction on *X* based on the information from *L* and learn the affinity matrix *W*, on which spectral clustering is performed to output the final result.

**Figure 1:**
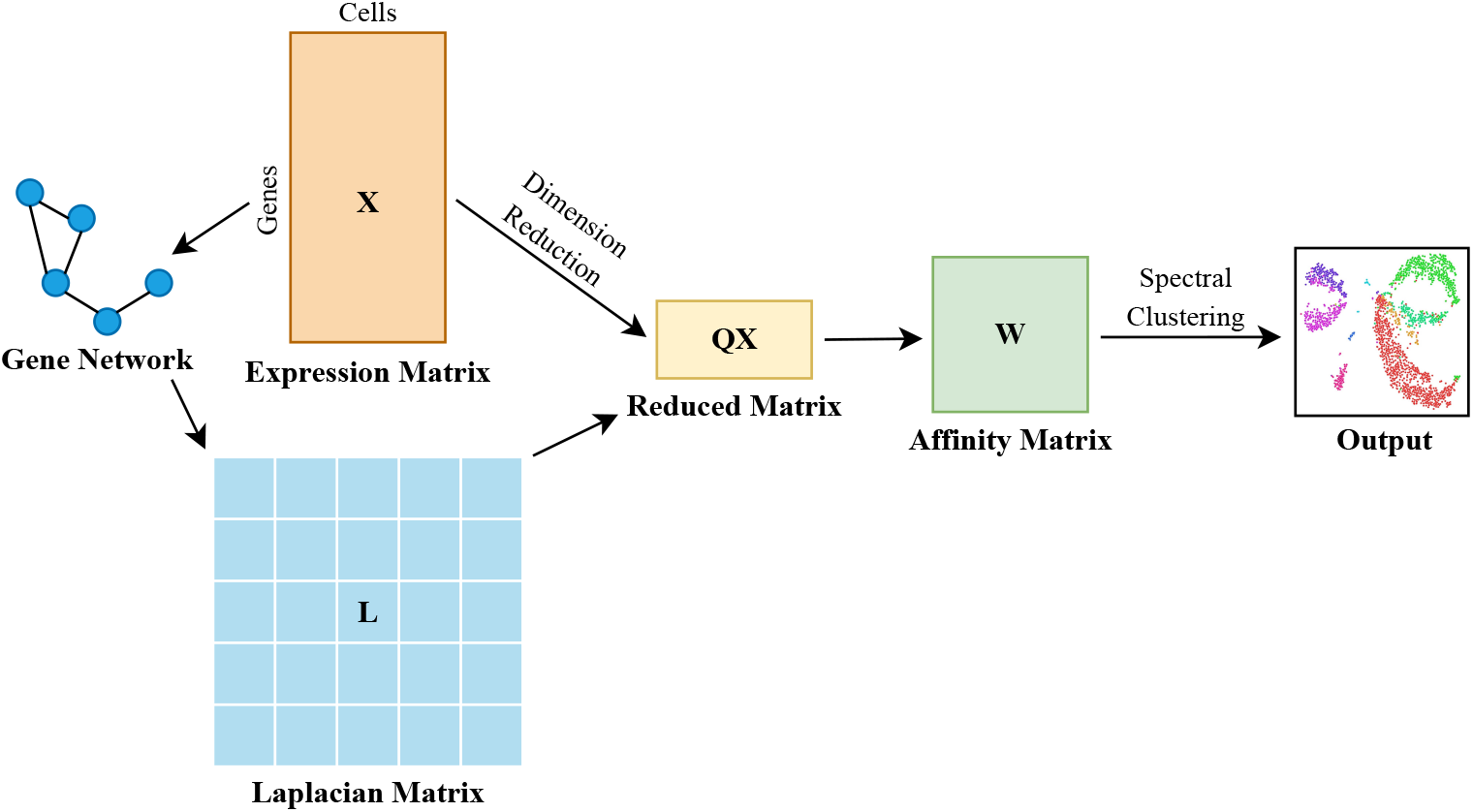
Overview of NetworkSSC pipeline

### 3.1 Gene Co-expression Network

Co-expression relationship exists between genes that are controlled by the same or functionally related transcriptional factors and thus have similar expression patterns (Weirauch, 2011). This relationship is often captured by an undirected graph named gene co-expression network. In such a network, each node represents a gene, and an edge connecting a pair of nodes indicates a significant co-expression relationship between the two genes (Stuart et al., 2003).

In a scRNA-seq data set, each row represents a specific gene. Since the transcript levels of two co-expressed genes often behave similarly across samples (Van Dam et al., 2018), the intrinsic co-expression relationship embedded in the data should be considered. However, traditional SSC does not take this prior information into account.

### 3.2 Design

We introduce a graph regularization term embedded with the Laplacian matrix of the gene network into the original cost function in SSC. According to the property of Laplacian matrix, this regularization term increases the weight of coefficients corresponding to intercluster samples and decreases the weight of coefficients corresponding to intra-cluster samples (He et al., 2017), and this modification effectively models the local geometric structure of the data and exploits the intrinsic relationship within the data. Besides graph regularization, we also perform dimension reduction on the expression matrix *X* beforehand for computational convenience. In our new formulation, the coefficient matrix *Z* is computed by

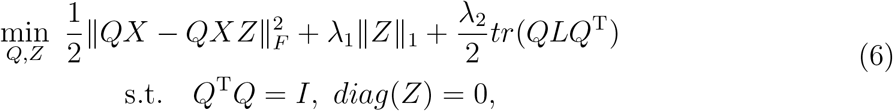

where *Q* ∈ ℝ^*d×p*^ is designed to reduce the dimension of *X* to *d*, and *L* ∈ ℝ^*p×p*^ is the gene network’s Laplacian matrix. The intrinsic relationship within the expression matrix *X* can be exploited in each iteration of optimization of *Q* and *Z*.

The successive steps are similar to those in SSC. Based on the optimal *Z*, we construct the affinity matrix *W*, on which we perform spectral clustering to obtain the final result.

### 3.3 Optimization

To solve *Q* and *Z* in the optimization problem in (6), we utilize a technique termed proximal alternating linearized minimization (PALM, Bolte et al., 2014). Our strategy is that in each iteration, we update *Q* and *Z* in an alternate manner.

Consider the optimization steps in the *t^th^* iteration. We first fix *Q*_*t*−1_ and solve *Z*, i.e.,

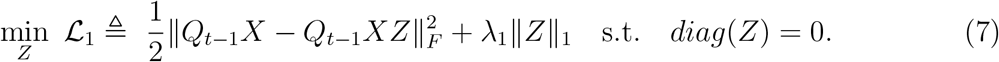

Let *μ*_*t*−1_ ≥ 1.01 · || (*Q*_*t*−1_*X*)^*T*^*Q*_*t*−1_*X*||_2_, then

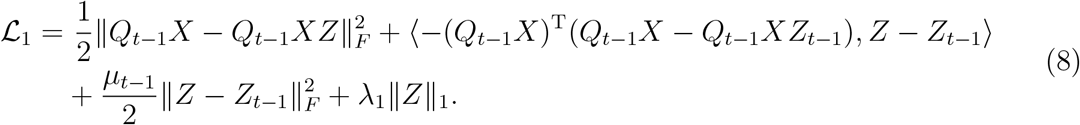

Let *G* ≜ −(*Q*_*t*−1_*X*)^*T*^(*Q*_*t*−1_*X*−*Q*_*t*−1_*XZ*_*t*−1_), then

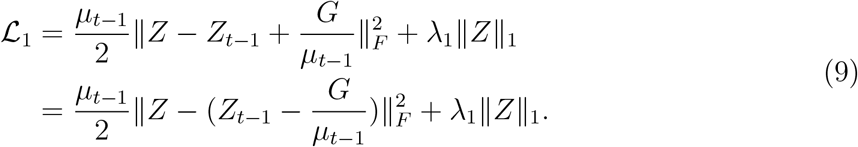

Let 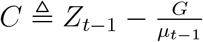, then we update *Z_t_* by

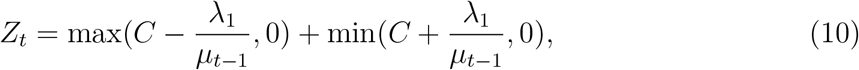

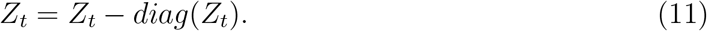

Then we fix *Z_t_* and update *Q_t_* by

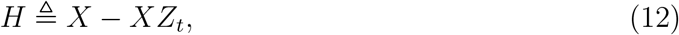

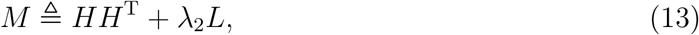

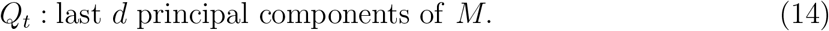

We also update *μ_t_* by

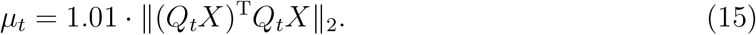

We continue this update process until the number of iterations exceeds an upper bound *kmax* or the update of *Z* and the cost function *f* in each step descend below a threshold value *ϵ*:

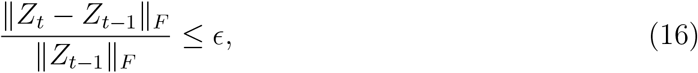

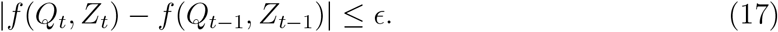

A possible value for *kmax* is 1000 and a possible value for *ϵ* is 0.001.

Finally, we consider the initialization of *Q* and *Z*. We initialize *Q* as

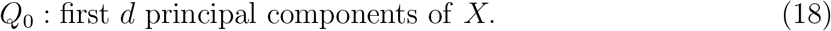

The initialization of *Z* can be found by solving

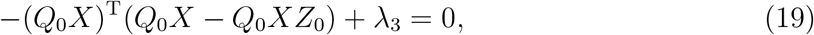

where we set *λ*_3_ = 1 as default. Thus,

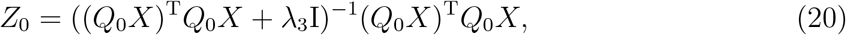

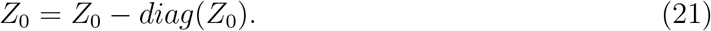

The complete procedures of NetworkSSC are summarized in Algorithm 1.

**Algorithm 1:**
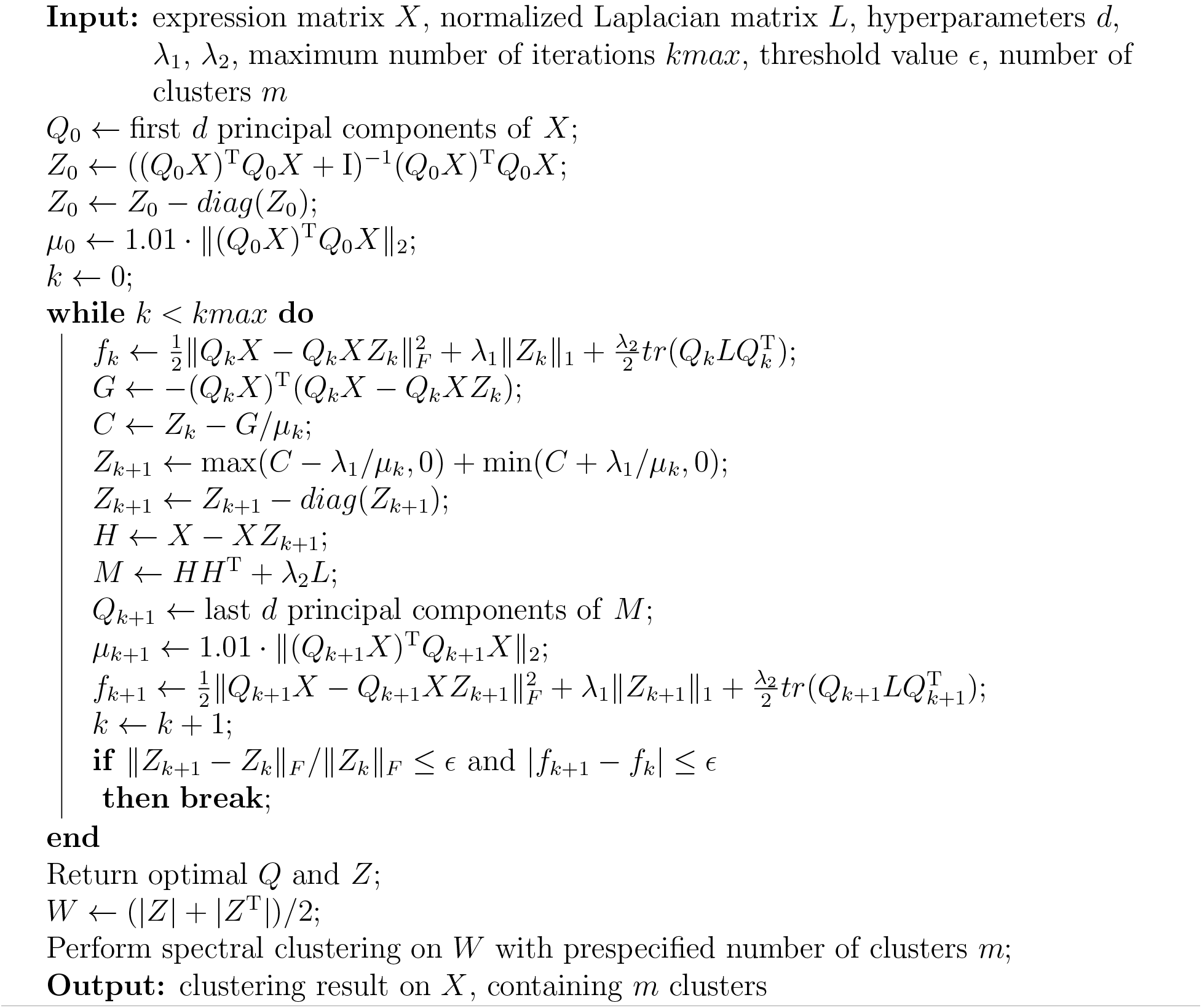
Network-guided sparse subspace clustering (NetworkSSC)

### 3.4 Treatment of Ultra-high Dimensional Data

In practice, when we apply NetworkSSC directly to real data, ultra-high dimensionality usually lead to unsatisfactory running speed and insufficient memory. In order to improve the computational efficiency, we design a shortcut to handle data sets with relatively large amount of genes, say, *p* > 5000. The idea is that, we equally partition the original data set into *m* blocks based on the *p* genes, where

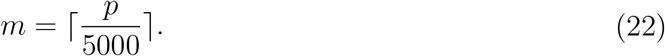

Note that the partition is completely at random. Then we run NetworkSSC on the *m* blocks separately, resulting in *m* affinity matrices *W*_1_, *W*_2_*, …, W_m_*. We average these matrices and obtain a single affinity matrix *W*, which can be regarded as having integrated all the information from the whole data. Finally, we perform spectral clustering on *W* to obtain the clustering result.

## 4 Data Experiments

### 4.1 Data Preparation and Pre-processing

We employ five scRNA-seq data sets of human organs (Baron et al., 2016; He et al., 2020; La Manno et al., 2016; Muraro et al., 2016; Segerstolpe et al., 2016). Our task is to assign the cells with different labels, each representing a specific cell type or subtype. All data sets have true labels of the cells, which can be used as ground truth to evaluate the clustering performance (Du`o et al., 2018). The basic features of the experimental data sets are summarized in Table 1.

**Table 1:** Basic features of experimental data sets

|  | Segerstolpe | Baron | Muraro | HeOrgan | Midbrain |
| --- | --- | --- | --- | --- | --- |
| Source Organ | Pancreas | Pancreas | Pancreas | Blood | Brain |
| Number of Genes | 2000 | 2000 | 19127 | 14552 | 19531 |
| Number of Cells | 702 | 2812 | 2126 | 1407 | 1695 |
| Number of Cell Types | 6 | 6 | 10 | 9 | 25 |

For each data set, we extract the corresponding gene network from a homosapiens gene network downloaded from HINT database (Das and Yu, 2012). For genes that do not match the full network, we still preserve them in the network and treat them as isolated points. Besides, we normalize the expression data for each gene to zero mean and unit variance.

### 4.2 Evaluation Metrics

To evaluate the performance of different clustering methods, we exploit two popular metrics: normalized mutual information (NMI, Pfitzner et al., 2009) and adjusted rand index (ARI, Hubert and Arabie, 1985). With *X* denoting the true labels and *Y* denoting the clustering labels, the definitions of NMI and ARI are as follows:

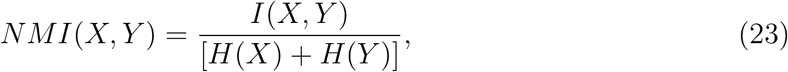

where *I* is the mutual information metric and *H* is the entropy.

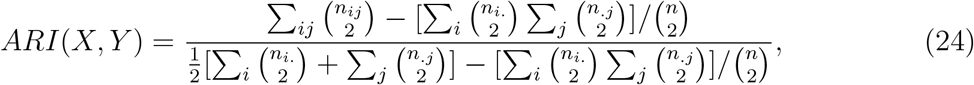

where *n_ij_* denotes the number of cells belonging to group *i* in true labels and group *j* in clustering labels, *n_i._* denotes the number of cells belonging to group *i* in true labels, and *n_.j_* denotes the number of cells belonging to group *j* in clustering labels. Both NMI and ARI can measure the consistency between the clustering results and the true labels.

### 4.3 Parameter Analysis

There are three hyperparameters to be set in NetworkSSC: the dimension of reduced data *d*, the *l*_1_ penalty factor *λ*_1_, and the Laplacian penalty factor *λ*_2_. An empirical rule for choosing *d* is to set it around 10% – 20% of the original dimension, so as improve the computational efficiency while retaining satisfactory clustering quality. In terms of *λ*_1_ and *λ*_2_, they are supposed to take relatively large values due to the dimension reduction procedure. In practice, we select the proper *λ*_1_ and *λ*_2_ by performing grid search within the range 100 – 1000.

### 4.4 Comparison Analysis of Clustering Methods

We test the proposed method on the selected data sets and compare the clustering result with five selected competitive methods, including traditional SSC, AdaptiveSSC, scDeepCluster, spectral clustering (SC), and K-Means. Traditional SSC is considered as the major target since NetworkSSC is a modification on it. The comparison results are summarized and visualized in Figure 2. Note that all tested methods have certain randomness due to the embedded K-Means procedure. In the experiment, we run each method 20 times and output the average NMI and ARI. The standard derivation of each method has been shrunk to negligible.

**Figure 2:**
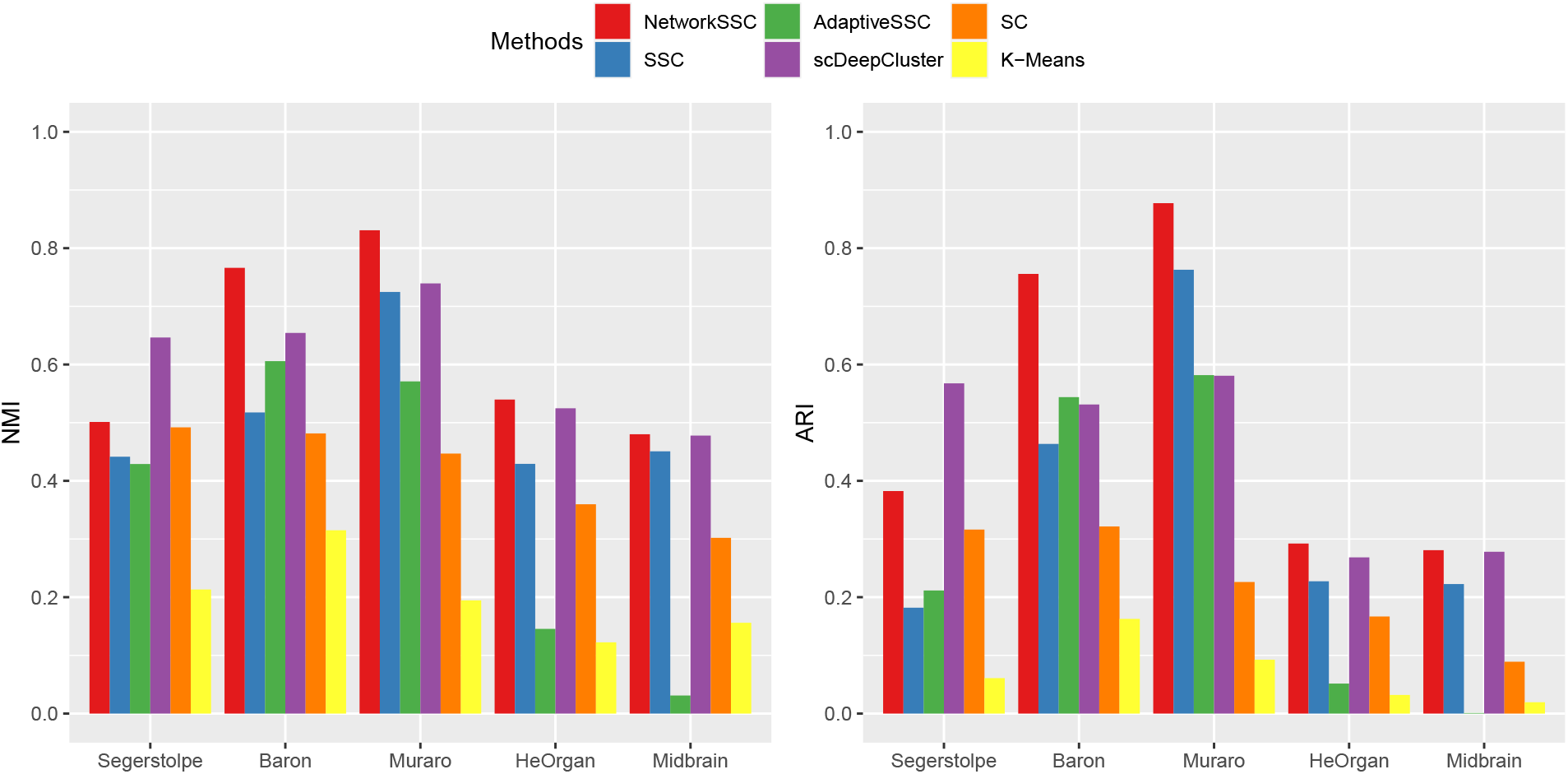
NMI and ARI of different methods on 5 data sets

According to Figure 2, NetworkSSC achieves the highest NMI and ARI over all competitive methods on most data sets. In specific, NetworkSSC outperforms traditional SSC on all data sets and with respect of both evaluation metrics, indicating that traditional SSC is significantly improved. Note that this dominance is robust against the dimensionality of these data sets, that is, whether the entire or a fraction of the sequenced genes are used to learn the cell type.

### 4.5 Robustness Analysis of Partition

In this section, we check whether employing different random partition will lead to significant variation on the clustering performance. Muraro and HeOrgan data are used as examples to illustrate the effect as they have relatively large amount of genes and partition is required. For each data set, we run NetworkSSC 10 times with different random partition. The box plots of NMI and ARI are shown in Figure 3. There are no outliers in the plot, which demonstrates that our algorithm is robust to the way of random partition on large data sets.

**Figure 3:**
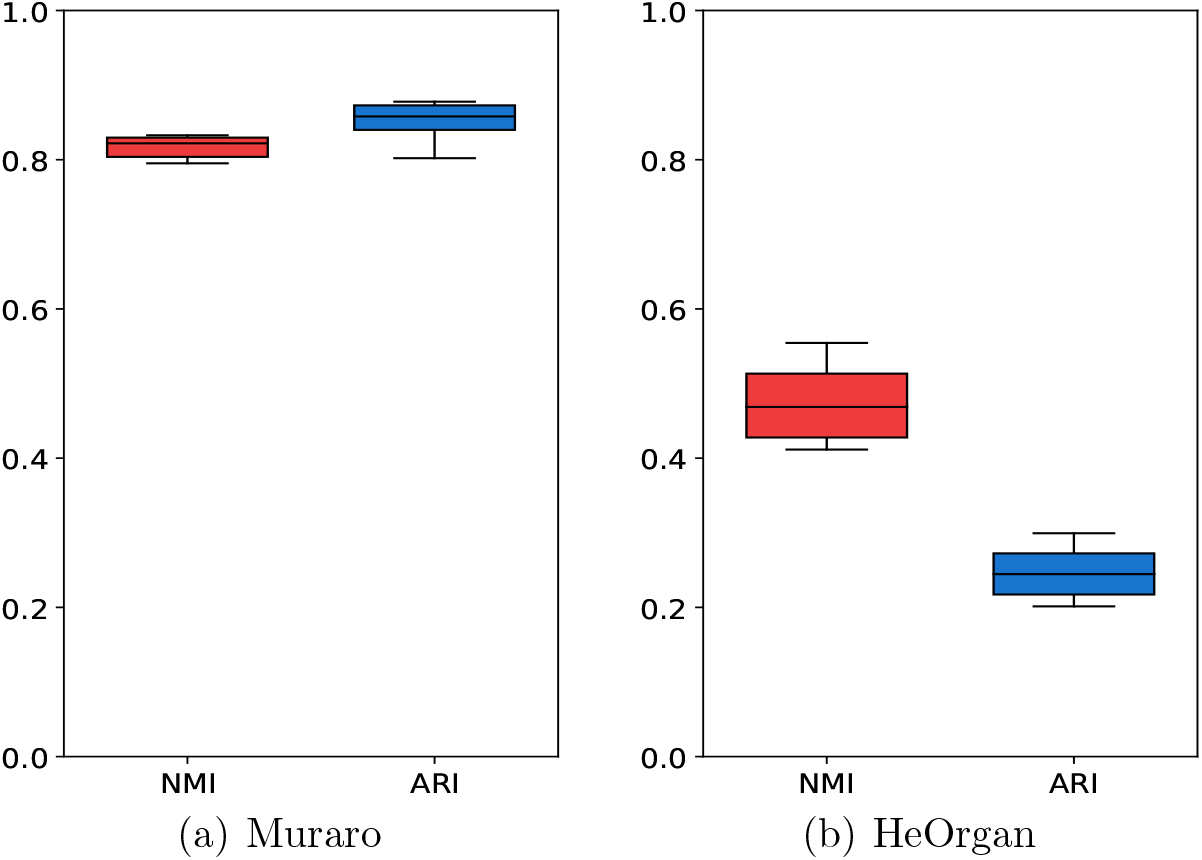
Effect of random partition on Muraro and HeOrgan data

## 5 Discussion and Conclusion

Cell type identification via unsupervised clustering is a fundamental issue in scRNA-seq data analysis. In this study, we propose a network-guided sparse subspace clustering approach called NetworkSSC. It assumes that the expression of cells within the same type lies in the same subspace. Furthermore, it introduces graph regularization to utilize the gene coexporession relationship for more effective cell clustering. Five real data sets are employed to evaluate the performance of NetworkSSC, on all of which NetworkSSC significantly improves the clustering accuracy compared with ordinary SSC. NetworkSSC also outperforms other four traditional and deep learning benchmarks on most data sets. The robustness analysis further validate the advantage and stability of our design.

Although we design a shortcut to handle large data sets and alleviate the computational burden, the computational efficiency of NetworkSSC is still unsatisfactory and should be improved in the future. Moreover, it is necessary to further refine our data processing procedures and adopt more state-of-art normalization methods to denoise the data. Attenuating the batch effect in large mixed data provides another promising possibility to improve our method and yield a higher accuracy of cell type identification.

## Data Availability

Data sets analyzed in this study can be found here: https://github.com/FrankYuan0916/NetworkSSC.

## Appendix 1: Comparison Results

**Table 2:**
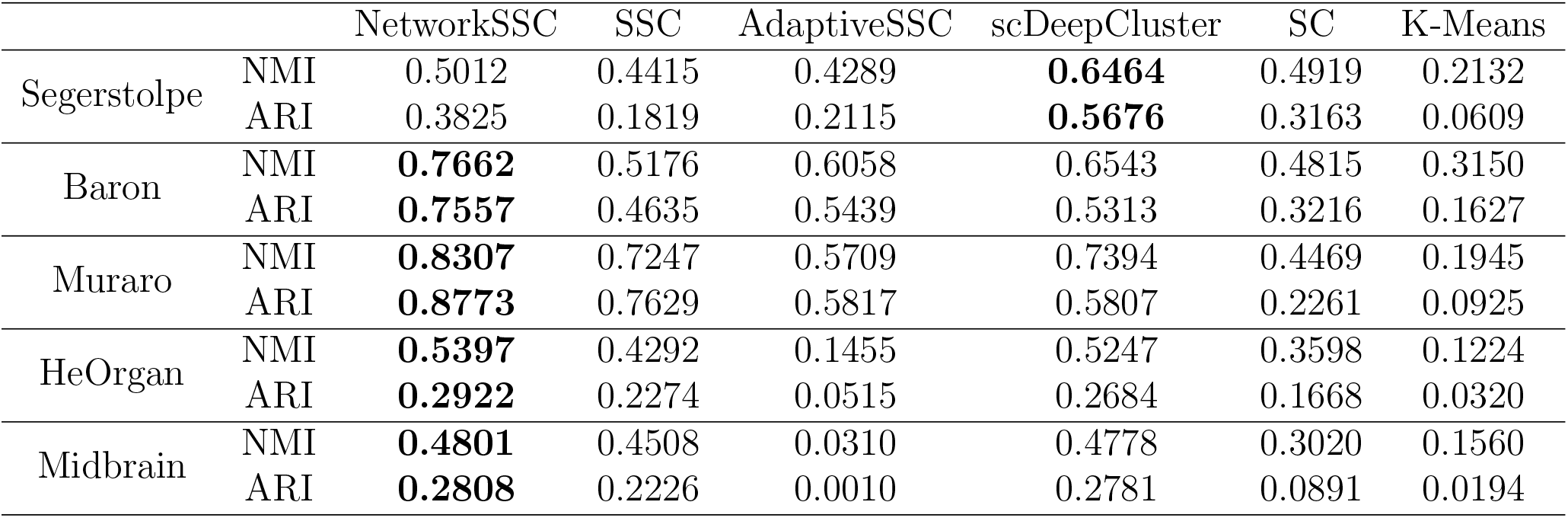
Comparison of NMI and ARI of different methods

## Notes

### Competing Interest Statement

The authors have declared no competing interest.

